# Electrospun cellulose acetate/gelatin nanofibrous wound dressing containing berberine for diabetic foot ulcer healing: *in vitro* and *in vivo* studies

**DOI:** 10.1101/787093

**Authors:** Hadi Samadian, Arian Ehterami, Saeed Farzamfar, Ahmad Vaez, Hossein Khastar, Mostafa Alam, Armin Ai, Zahra Allahyari, Majid Salehi

**Author notes:** Arian Ehterami, Ph.D. student. Saeed Farzamfar, Ph.D. student. Mostafa Alam, DMD, Postgraduate student. Armin Ai, Dental student. Majid Salehi, Ph.D. Corresponding author.

## Abstract

Functional dressing with tailored physicochemical and biological properties is vital for diabetic foot ulcer (DFU) treatment. Our main objective in the current study was to fabricate Cellulose Acetate/Gelatin (CA/Gel) electrospun nanofibrous mat loaded with berberine (Beri) as the DFU dressing. The results demonstrated that the diameter of the nanofibers was around 502 nm, the tensile strength, contact angle, porosity, water vapor permeability, and water uptake ratio of CA/Gel nanofibers were around 2.83 MPa, 58.07, 78.17 %, 11.23 mg/cm2 hr, and 12.78 respectively, while these values for CA/Gel/Beri nanofibers were 2.69 ± 0.05 MPa, 56.93 ± 1, 76.17 ± 0.76 %, 10.17 ± 0.21 mg/cm2 hr, 14.37 ± 0.42 respectively. The bacterial evaluations demonstrated that the dressings are an excellent barrier against bacterial penetration with potent antibacterial activity. The animal studies depicted that the collagen density and angiogenesis score in the CA/Gel/Beri treated group were 88.8±6.7 % and 19.8±3.8, respectively. These findings implied that the incorporation of berberine did not compromise the physical properties of dressing, while improving the biological activates. In conclusion, our findings implied that the prepared mat is a proper wound dressing for DFU management and treatment.

## 1. Introduction

Diabetes mellitus is classified in metabolic disease which has various complications such as chronic wounds, arterial damage, and neuropathy in the result of uncontrolled blood sugar. The wound healing process is a complex and multiphase process which is delayed in diabetic patients because of various complexities [1, 2]. In these patients, the angiogenesis and re-epithelialization are poor because of low interaction between growth factors and their target site. The severe inflammation is another deleterious factor which is because of neutrophil infiltration. Moreover, diabetic foot ulcer (DFU) is another complication which is the consequence of intense inflammation, limited nutrients, and poor circulation. Despite the tremendous breakthroughs last decades, the effective treatment of diabetic foot ulcer remains a challenge [3].

Since DFU is an acute wound, it needs to be dressed with proper dressing materials able to enhance the healing process and also isolate the wound site from pathogen microorganism [4]. Moreover, the wound dressing must be able to absorb the exerted exudates from the wound and also provide an optimum moist environment to expedite the healing process. In addition, the healing process of diabetic foot ulcer required to be accelerated via bioactive molecules or drugs and the proposed dressing must be able to load a proper amount of the drug and release it in a sustain release manner [5-7]. A wide range of biomaterials and nanostructured materials have been evaluated as the wound dressings for DFU such as natural and synthetic polymers in the forms of hydrocolloids, hydrogels, foams, and electrospun nanofiber dressing [8, 9].

Electrospun nanofibrous dressings offer a wide range of promising possibilities suitable for wound dressing applications. Electrospinning is a sophisticated and efficient technique which provides a low-cost, scalable, flexible, relatively simple approach for nanofibers fabrication from a very wide variety of synthetic and natural substance [10-13]. The porosity and the pore size of the electrospun mats can be adjusted to inhibit microorganism penetration, while oxygen can easily pass through the dressing and reach to the wound site. Interestingly, the water vapor transmission can be tailor to provide the ideal moisture for the wound healing process. The high surface area of nanofibers is favorable for drug loading and sustained deliver. The interested drugs, natural substance or bioactive molecules can be adsorbed onto the surface of nanofibers or encapsulated into the nanofibers matrix. Moreover, the electrospun nanofibrous dressings are self-standing and their handling during the wound treatment is easy [14, 15].

Electrospun nanofibers have been fabricated from variety of natural and synthetic polymers and applied as the wound dressing. Among them, natural polymers have grabbed considerable attention due to their fascinating properties. Most of them are biocompatible, non-toxic, biodegradable, abundant, inexpensive sources, diverse, renewable, and versatile [16, 17]. Cellulose is one of the most abundant natural polymers on earth which has various derivative, but the most fascinating derivative is cellulose acetate (CA), the acetate ester form of cellulose [18-20]. CA is applicable in various applications such as membrane separation, biomedical, textile fibers, cigarette industries, and plastics [21-24]. The biomedical applications of CA are mainly categorized in drug delivery systems, tissue engineering, and wound dressing [25-28]. Biocompatibility, water absorption abilities, and good interaction with fibroblast cells have mad CA a good candidate for wound dressing applications [29]. Gelatin is a widely used natural polymer with fascinating biological properties such as biocompatibility, biodegradability, and bioactivity which is obtained from collagen hydrolysis [30-32]. Various forms of gelatin can be fabricated such as the hydrogel, layered, freeze-dried, and micro- and nanofibers for different applications. Due to the presence of cell adhesion domains such as Arg-Gly-Asp (RGD) domain in the structure of gelatin, it is widely used in tissue engineering and wound dressing applications [33-35].

In addition to the structural requirements, a proper DFU dressing should have an active ingredient to either enhance the healing process or even provides the antibacterial property. Various types of biological, natural, and chemical moieties have used to induce biological functions to the dressings. Berberine is a natural substance belongs to the alkaloid family found in the rhizome, roots, and stems of various plants such as Oregon grape, Goldenseal, and Barberry [36]. Berberine is known for its anti-diabetic, antimicrobial, and anti-inflammatory activates. Moreover, some studies reported the diabetic wound healing efficacy of berberine [37]. Accordingly, the main objective of our study is to fabricate CA/Gel electrospun nanofibrous mat containing berberine as a diabetic foot ulcer wound dressing.

## 2. Materials and methods

### 2.1. Chemicals

All the chemicals and materials were, unless otherwise noted, purchased from Sigma-Aldrich (St. Louis, USA), Thermo Fisher Scientific (Waltham, MA, USA), and Merck (Darmstadt, Germany).

### 2.2. Fabrication of nanofibrous wound dressing

A proper amount of gelatin (bovine skin, type B) and CA [Mw= 30 kDa, acetyl content = 39.70% (w/w)] were separately dissolved in 1,1,1,3,3,3-hexafluoro-2-Propanol (HFP) to obtain the final concentration of 6% (w/v). CA (6% (w/v)) and Gel (6% (w/v)) were combined at the weight ratio of 25:75 and stirred for 24h. In the next step, the berberine was added to the resulted solution in the amounts of 1% (w/v) and stirred for 24 h. The commercial electrospinning apparatus (NanoAzma, Tehran, Iran) was used to fabricate the electrospun wound dressing. Briefly, the CA/Gel/Beri solution was loaded into a 10 mL disposable syringe ending to a 25-gauge stainless steel blunted needle. The needle was connected to the power supply and the polymer solution was pumped onto the tip of the needle via a syringe pump at the constant rate. The applied voltage, the feeding rate, and the nozzle to the collector distance were 15 kV, 0.2 mL/h, and 15 cm, respectively. The fabricated mats were incubated with the vapor of 10 % (w/v) glutaraldehyde (GA) for 16 h to induce cross-linking between the nanofibers. The cross-linked nanofibers then washed thoroughly by distilled water to remove unreacted GA followed by drying at room temperature.

### 2.3. Characterization of the wound dressing

#### 2.3.1 Scanning Electron Microscopy analysis

The produced wound dressings were sputter coated with a thin layer of gold using a sputter coater (KYKY Technology Development, Beijing, China) and imaged using a scanning electron microscope (SEM; KYKY Technology Development, Beijing, China) at an accelerating voltage of 26 kV.

#### 2.3.2. Mechanical strength measurement

The tensile strength of the fabricated wound dressings was measured by a uniaxial tensile testing device (Santam, Karaj, Iran) with an extension rate of 1 mm/min according to ISO 5270:1999 standard test methods.

#### 2.3.3. Contact angle measurement

Water contact was measured as the indication of hydrophilicity/hydrophobicity nature of the dressings via a static contact angle measuring device (KRUSS, Hamburg, Germany). The water droplet was poured onto the three different points of each sample and the resulted angle between the water droplet and the surface of each specimen was averaged and reported.

#### 2.3.4. Water vapour permeability (WVP) test

The flexible bottles permeation test (Systech, UK) was used to measure water vapor permeability (WVP) of the fabricated dressings. The dressings-capped bottles were incubated at 33 °C for 12 hours, the evaporated water through the dressing was measured, and equation 1 was used to calculate WVP at steady-state.

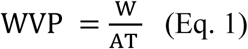

Where W, A, and T stand for the mass of water lost, area (1.18 cm^2^) of dressing, and exposure time, respectively.

#### 2.3.5. Water uptake ratio evaluation

The water uptake ability of the mats was measured based on our previous studies [38, 39]. The dry dressing was weighted (W_0_), immersed in distilled water at room temperature for 24 h, and then the wet samples were weighted again (W_1_). Equation 2 was used to calculate the water-uptake capacity of the dressings. Three measurements were conducted for each specimen and the averages were reported.

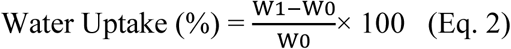

#### 2.3.6. Porosity assessment

The porosity of the electrospun dressings was measured based on the liquid displacement technique using equation 3 [40]. Briefly, the mat of weight W was immersed in a graduated cylinder containing a known volume (V1) of ethanol and the resulted volume then was recorded as V2. After 10 min the samples were removed and the residual ethanol volume recorded as V3.

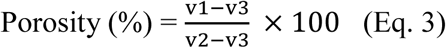

#### 2.3.7. Weight loss measurement

Weight loos of the dressings was measured as the function of degradation based on our previous study [38]. Briefly, proper amount of dressings were weighted (W_0_), incubated in PBS solution, extracted for the solution after 7 and 14 days, and precisely weighted (W_1_). Equation 4 was used to calculate the weight loss.

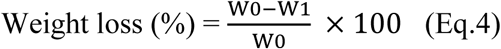

### 2.4. Microbial penetration test

The resistance of the fabricated dressings against microbial penetration was assessed by the microbial penetration test. Briefly, 10 ml vials (test area: 0.8 cm^2^) containing 5ml of Brain heart infusion (BHI) broth culture medium (Merck, Germany) was capped with the prepared mates, kept at ambient conditions, and growth of the bacteria into the culture medium was measured after 3 and 7days. Bottles covered with the cotton ball and open vials served as negative and positive controls, respectively. The turbidity as the indication of microbial contamination was measured by spectroscopy approach at 600 nm using a microplate spectrophotometer (n=3).

### 2.5. Antibacterial growth assay

The antibacterial activities of the prepared nanofibers were conducted based on the time-kill assay against *Staphylococcus aureus* and *Pseudomonas aeruginosa* [41]. Briefly, the prepared CA/Gel nanofibers were added to bacterial suspensions (previously adjusted to 1×10^7^ CFU/ml) at concentrations of 1/2-fold of the Minimum Inhibition Concentrations (MIC). 0.5 mL of each suspension was incubated at 37 °C with gentle agitation in a shaking water bath for 24 h. After the incubation period, the suspension (10 μL) was serially-diluted and inoculated on agar plates and incubated for 1, 2, 4, and 24 h in aerobic incubation condition at 37 °C. Then, the number of viable bacteria colonies was counted.

### 2.6. Blood compatibility or hemolysis assay

The hemolysis assay was conducted on human whole blood anticoagulated and diluted with normal saline. The samples were incubated with 200 µl of blood samples at 37 °C for 60 min followed by centrifugation at 1500 rpm for 10 min. Then the absorbance of the resulted supernatant was read at 545 nm using a Multi-Mode Microplate Reader (BioTek Synergy™ 2). The negative control and positive control were the whole blood diluted in normal saline and the whole blood diluted in deionized water, respectively. Equation 4 was used to calculate the hemolysis percent [42].

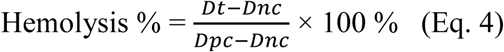

where Dt, Dnc, and Dpc is the absorbance of the sample, the absorbance of the negative control, and absorbance of the positive control, respectively.

### 2.7. Cell culture studies

L929 murine fibroblastic cell line was obtained from Pasteur Institute, Tehran, Iran and used to evaluate the biocompatibility of the prepared dressing. Dulbecco’s modified Eagle’s medium: nutrient mixture F-12 (DMEM/F12; Gibco, Grand Island, USA) enriched with 10% (v/v) fetal bovine serum (FBS; Gibco, Grand Island, USA), 100 unit/mL of penicillin, and 100 µg/mL of streptomycin. The dressings were cut spherically and put into the bottom of each well of a 96-well plate followed by UV light exposure for 1 h for sterilization. Then, the nanofibrous mat was washed with PBS twice and once with DMEM/F12 and then seeded with 1×10 ^4^ cells. 3-(4, 5-Dimethylthiazol-2-yl)-2, 5-Diphenyltetrazolium Bromide (MTT) assay kit was used to evaluate the proliferation of the seeded cells based on the previously described method [43]. The positive control was the wells without dressings, the experiment was done triplicated, and the average data reported.

The morphology of cultured cells in the fabricated nanofibers was observed using SEM imaging. Briefly, after 24 h the seeded cells were fixed in 4 % paraformaldehyde for 1 h at room temperature, dehydrated in graded ethanol, sputter coated with a thin layer of gold, and observed at an accelerating voltage of 26 kV.

### 2.8. Animal studies

#### 2.8.1. Induction and assessment of diabetes

The animal studies were conducted on male adult Wistar rats purchased from Pasteur Institute, Tehran, Iran. Diabetes was induced by intraperitoneal injection of a single dose of 55 mg/kg streptozotocin (STZ) in citrate buffer (pH 4.4, 0.1 M). The equal volume of citrate buffer was injected intraperitoneally to the age-matched control rats. Retro-orbital plexus technique was used to collect the blood samples and GOD-POD (glucose oxidase-peroxidase) diagnostic kit (Accurex Biomedical Pvt. Ltd.,Mumbai,India) was used to measure serum glucose levels and the levels more than 250 mg/dL confirmed the diabetes induction [44].

#### 2.8.2. In vivo wound healing study

The animal study was conducted on 24 male adult Wistar rats based on the instruction of the ethics committee of Shahroud University of Medical Sciences. General anesthesia was induced by ip injection of Ketamine 5% / Xylazine 2% (70 mg ketamine and 6 mg Xylazine /1 kg body weight). The full-thickness excisional wound model was created on foot of each rat as a rectangular pattern marked on the dorsal surface of the foot using a flexible transparent plastic template, and then a layer of skin in full thickness with a standard area of 2 mm × 5 mm was removed. The animals were divided into 4 groups (6 rats per group) and the wounds were treated with the CA/Gel, CA/Gel/Beri, and the sterile gauze as the negative control.

After 16 days post-surgery, the animals were sacrificed by ketamine overdose injection and the wound tissue was harvested and fixed in 10% buffered formalin. The harvested tissues were processed, embedded in paraffin, sectioned, and stained with hematoxylin-eosin (H&E) and Masson’s trichrome (MT). An independent pathologist observed and interpreted the prepared slides under a light microscope (Carl Zeiss, Thornwood, USA) with a digital camera (Olympus, Tokyo, Japan) and photographed at ×100, and ×200 magnification. Epithelialization, inflammatory cell infiltration, fibroplasia, and granulation tissue formation assessed in different groups, comparatively.

Histomorphometry analysis was conducted at 16 days post-treatment to further evaluate the healing process. The assessment was semi-quantitatively by comparative analysis on 5 point scales: 0 (without new epithelialization), 1 (25%), 2 (50%), 3 (75%), and 4 (100%). Moreover, collagen density as informative data was measured and analyzed on the wound site using computer software Image-Pro Plus® V.6 (Media Cybernetics, Inc., Silver Spring, USA).

### 2.9. Statistical Analysis

Minitab 17 software (Minitab Inc., State College, USA) was used to statistically analyse the obtained data Student’s *t*-test and the data were expressed as the mean ± standard deviation (SD). In all evaluations, *P* < 0.05 was considered as the statistically significant.

## 3. Results and discussion

### 3.1 Characterization results

#### 3.1.1. The morphology of nanofibers

Various characterization methods were used to assess the properties of the fabricated nanofibers. The morphology of the prepared nanofibers was observed by using SEM imaging (Fig. 1). The SEM image showed that the fabricated CA/Gel/Beri nanofibers are uniform and straight without any beads and deformities. The image analysis using ImageJ software (National Institutes of Health, Bethesda, USA) showed that the diameter of the nanofibers was 502 ± 150 nm.

**Fig. 1.**
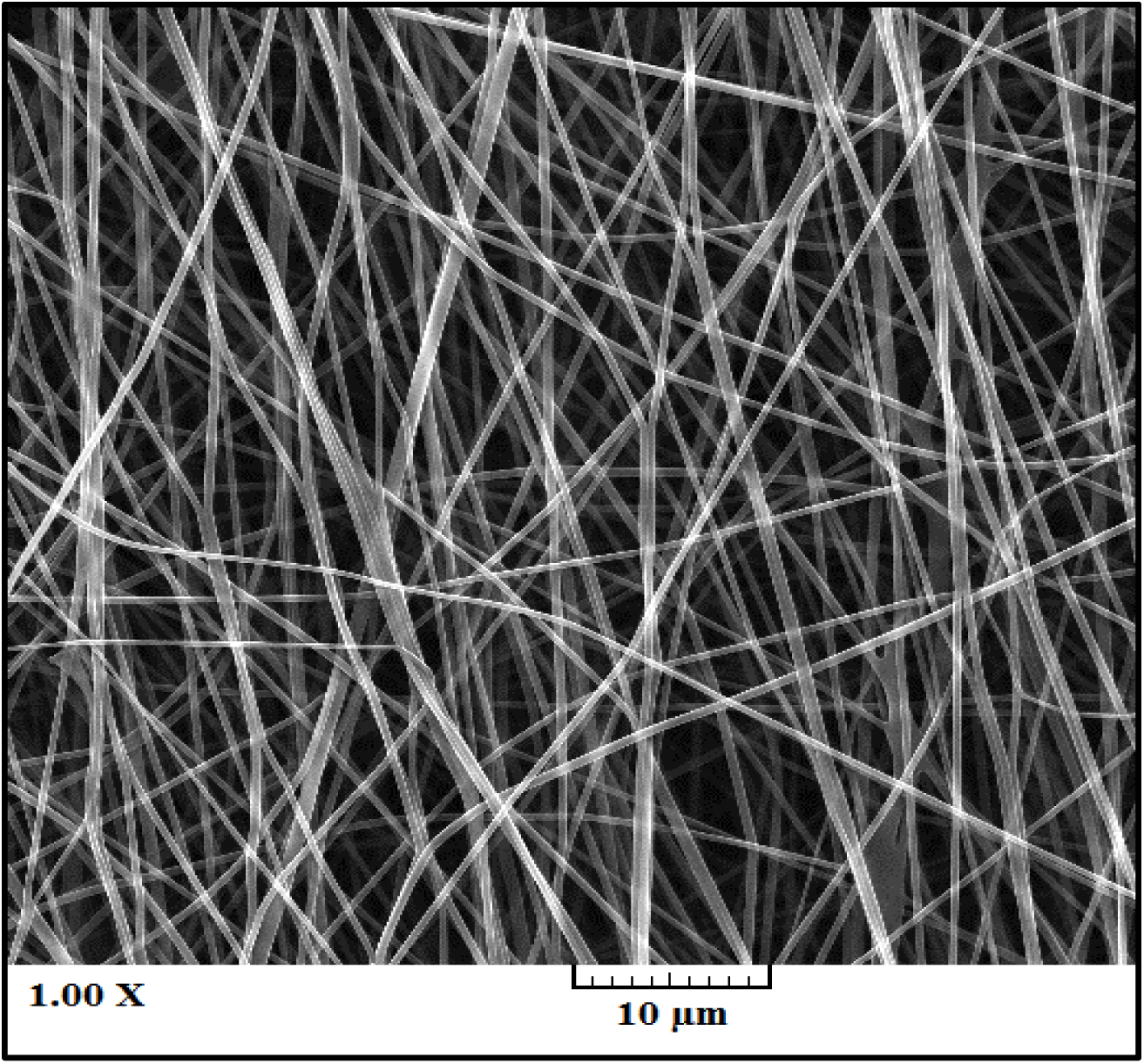
SEM image of the electrospun CA/Gel/Beri nanofibers.

#### 3.1.2. Mechanical property

Mechanical properties of the nanofibers were measured by tensile strength method based on ISO 5270:1999 standard test methods. The results showed that the incorporation of berberine compromised the mechanical property and reduced the tensile strength from 2.83 ± 0.08 to 2.69 ± 0.05 MPa. Our previous study [39] showed that the addition of taurine to electrospun Poly (ε-caprolactone)/Gelatin nanofibers reduced the mechanical property. This can be attributed to the berberine-induced reduction in the physical interaction between the polymer chains. Berberine weakened interchains and intrachain physical interaction and forces such as van der Waals forces and hydrogen bonding.

#### 3.1.3. Porosity

The liquid displacement technique was used to measure the porosity of the fabricated CA/Gel/Beri nanofibers and the results showed that the porosity of CA/Gel and CA/Gel/Beri nanofibers were 78.17 ± 1.04 and 76.17 ± 0.76 %, respectively. However, the porosity of the fabricated CA/Gel/Beri nanofibers are in the acceptable range, Water vapor permeability (WVP) should be measured along to conclude the efficacy of the prepared CA/Gel/Beri nanofibers.

#### 3.1.4. Wettability

A suitable dressing should be able to absorb the wound exudates and maintain the moisture of the wound site. These criteria are under the influence of the surface wettability and hydrophilicity of the dressing. The wettability of the fabricated dressings was measured based on water contact angle method and the results are presented in Table 1. The water contact angle of CA/Gel and CA/Gel/Beri nanofibers were 58.07 ± 2.35° and 56.93 ± 1°, respectively, which indicated that the fabricated dressings are hydrophilic and proper for absorbing exudates and maintaining the moisture of the wound bed. Liu et al [45] demonstrated that the incorporation of gelatin can increase the surface wettability of cellulose acetate due to its hydrophilic nature.

**Table 1.**
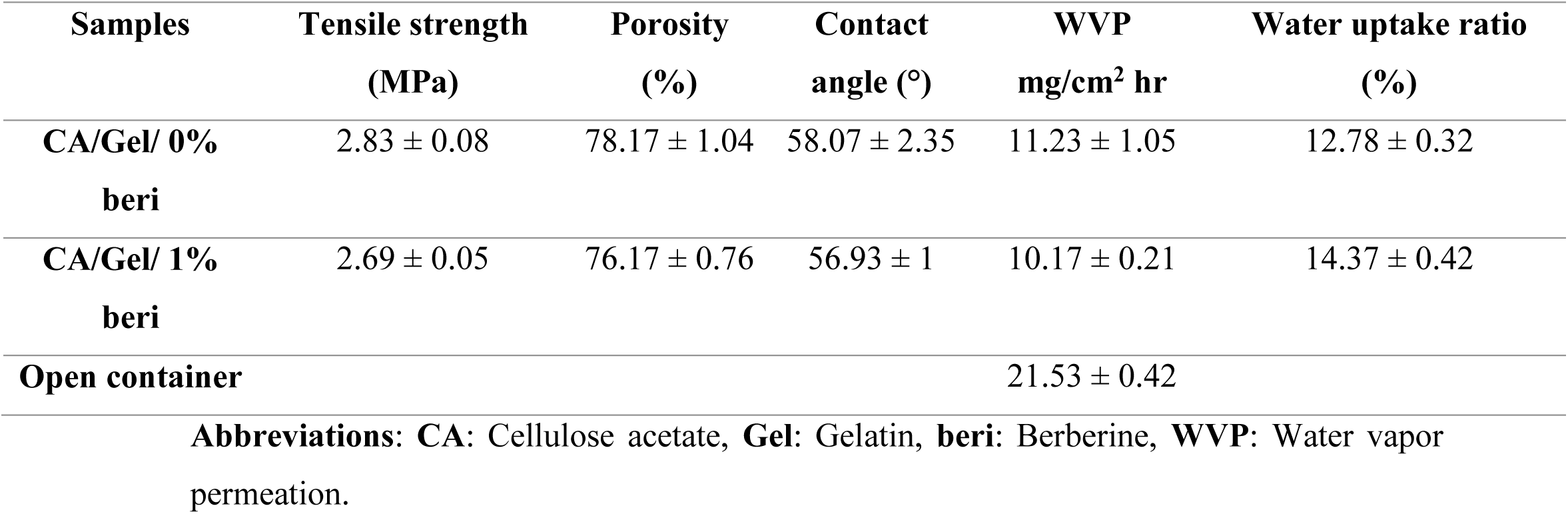
Characteristic of the fabricated CA/Gel/Beri nanofibers.

| Samples | Tensile strength<br>(MPa) | Porosity<br>(%) | Contact<br>angle (°) | WVP<br>mg/cm<sup>2</sup> hr | Water uptake ratio<br>(%) |
| --- | --- | --- | --- | --- | --- |
| CA/Gel/ 0%<br>beri | 2.83 ± 0.08 | 78.17 ± 1.04 | 58.07 ± 2.35 | 11.23 ± 1.05 | 12.78 ± 0.32 |
| CA/Gel/ 1%<br>beri | 2.69 ± 0.05 | 76.17 ± 0.76 | 56.93 ± 1 | 10.17 ± 0.21 | 14.37 ± 0.42 |
| Open container |  |  |  | 21.53 ± 0.42 |  |

**Abbreviations:** CA: Cellulose acetate, Gel: Gelatin, beri: Berberine, WVP: Water vapor permeation.

#### 3.1.5. Water vapor permeability and water-uptake capacity

The vapor exchange through the dressing is a critical property determining the efficacy of the dressing. High WVP value dehydrates the wound and induces scar formation, while the low WVP value delays the wound healing process due to the deposited exudates. Therefore, a proper dressing should exhibit an optimum value of WVP. The results showed that the WVP value for CA/Gel dressing was 11.23 ± 1.05 mg/cm2 hr, while the addition of berberine reduced to 10.17 ± 0.21 mg/cm2 hr which both values are significantly lower than the control group (the open container).

As shown in Table 1, the water uptake ratio of CA/Gel was 12.78 ± 0.32 % and the incorporation of berberine increased the ratio to 14.37 ± 0.42 %. This enhancement in the water uptake ratio can be related to the hydrophilic nature of berberine. These results demonstrate that the fabricated CA/Gel and CA/Gel/Beri can properly absorb the wound exudates and subsequently improve the wound healing process.

#### 3.1.6. Weight loss assay findings

The degradation rate of the prepared CA/Gel and CA/Gel/Beri was measured in PBS at days 7 and 14 (Fig. 2). The results represented that the fabricated dressings were not degradable during 14 days and the highest weight loss, around 80 %, was observed in CA/Gel/Beri group at days 14.

**Fig. 2.**
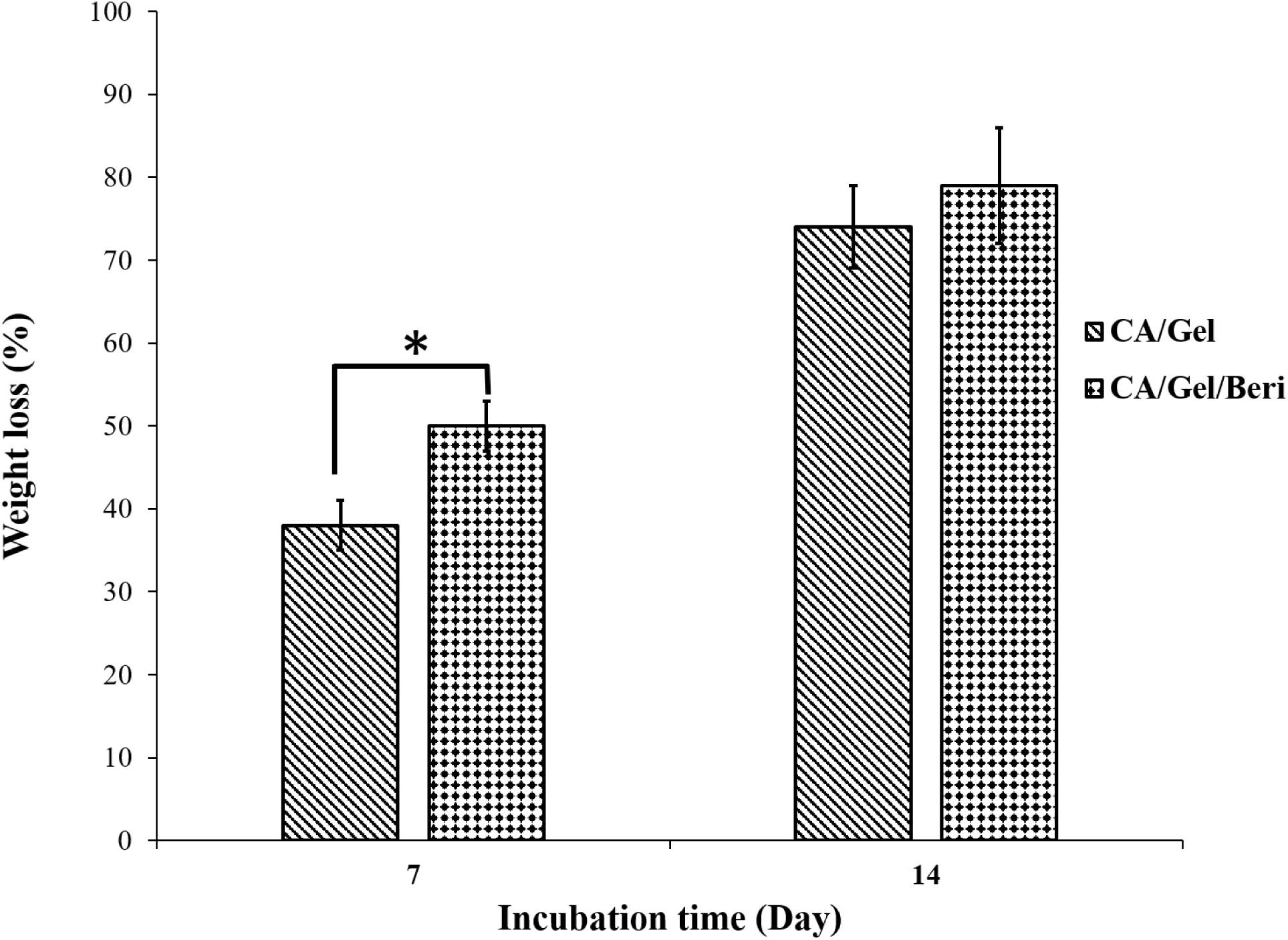
Weight loss percentage of the fabricated dressings at different time points (7 and 14 days). Values represent the mean ± SD, n = 3, *p < 0.05

As shown in Fig. 2, the incorporation of berberine accelerated the weight loss of CA/Gel nanofibers at both time intervals which was statistically significant at 7 days (*p < 0.05). The observed increased weight loss can be related to the hydrophilic nature of berberine which enhanced the interactions between CA/Gel and water molecules. Moreover, berberine may reduce the physical interactions between the polymer chains which facilitates the degradation rate. However, in the wound dressing applications there is no need for biodegradability.

### 3.2. Microbial evaluations findings

#### 3.2.1. Microbial penetration

A proper wound dressing must withstand against microbial invasion through the dressing and show acceptable microbial barrier property. In this experiment, the negative control was the tube closed with a cotton ball to test the sterilization procedure, while the positive control was open tube and tested to ensure the growth possibility of bacteria in the nutrient broth.

As shown in Fig. 3, the fabricated dressings prevent bacterial transition and negligible colonies were grown in the culture media which was statistically significant compared with the positive control (P<0.005). Moreover, the cloudiness of the nutrient broth was further evaluated by Spectrophotometer at 600 nm and the results are shown in Fig. 4.

**Fig. 3.**
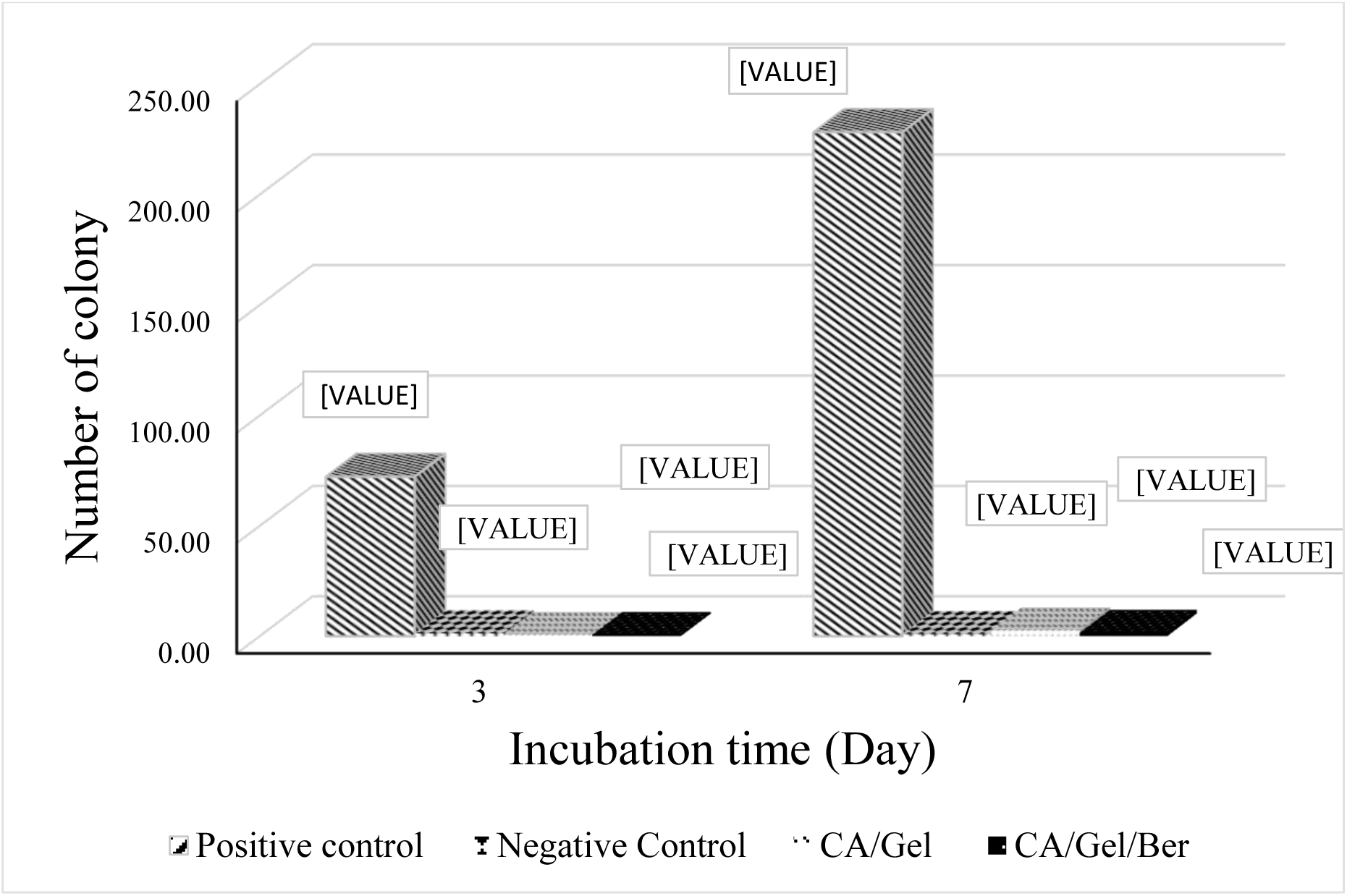
Microbial barrier property of the fabricated dressing after 3 and 7 days incubation, measured by colony counting assay. Values represent the mean ± SD, n = 3.

**Fig. 4.**
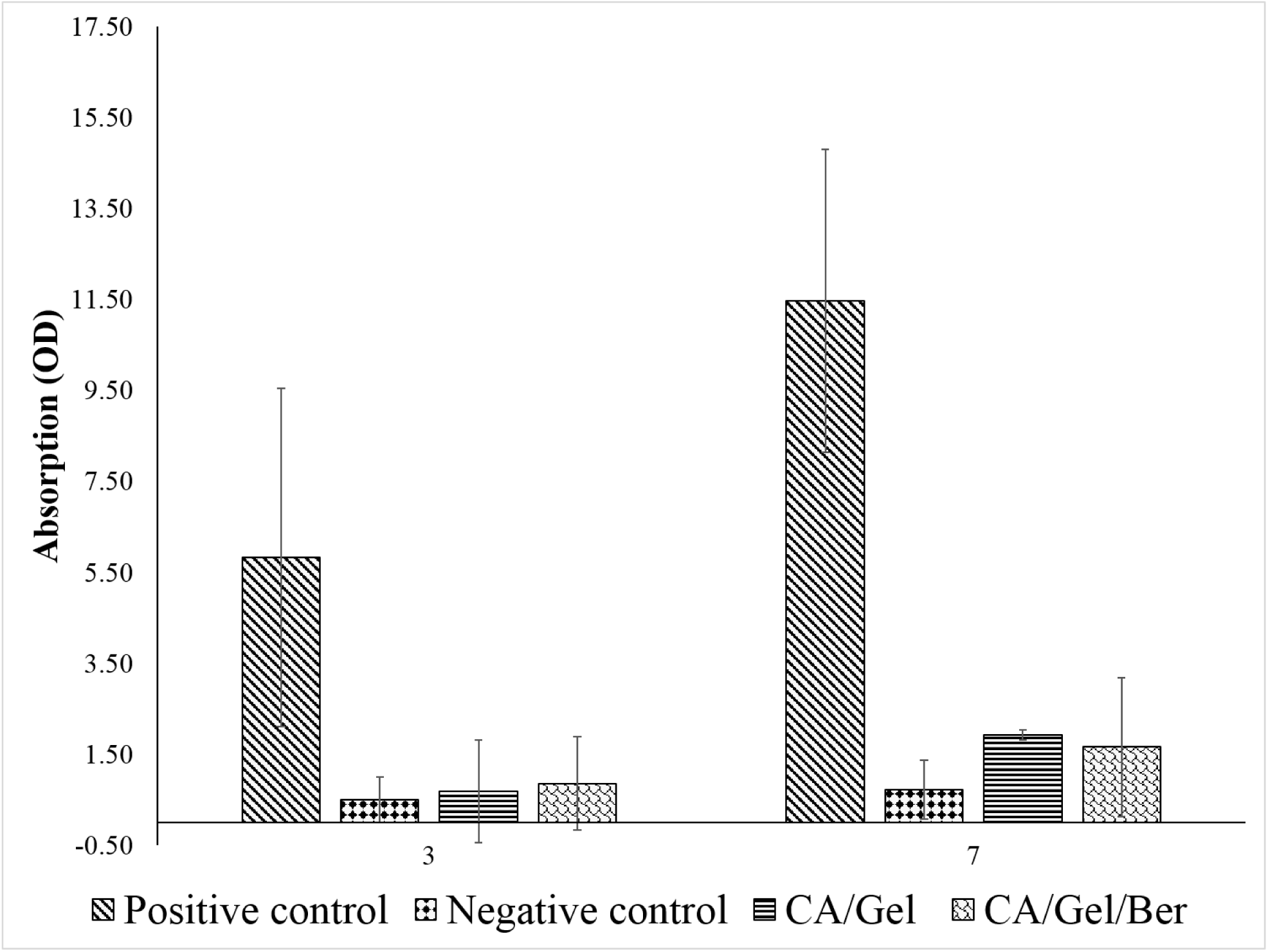
Microbial barrier property of the fabricated dressing after 3 and 7 days incubation measured by Spectrophotometer at 600 nm. Values represent the mean ± SD, n = 3.

The turbidimetry assay results confirmed the colony counting assay findings which indicated that the fabricated dressings exhibited excellent microbial barrier property. The results demonstrated that the microbial penetration of CA/Gel/Beri dressing was lower than CA/Gel which can be attributed to the antibacterial property of berberine. The antibacterial activity of berberine was shown by Peng et al. [46]. Moreover, it is shown that even 64 layers of gauze were not able to stand against bacterial penetration into the wound [47]. Hence, these results imply that the fabricated dressings are suitable for wound care application.

#### 3.2.2. Antibacterial assay findings

An effective wound dressing should possess the antibacterial activity along with bacterial penetration barrier. The antibacterial activities of the fabricated dressings were assessed by time-kill assay against gram-positive and gram-negative bacterium *Staphylococcus aureus* and *Pseudomonas aeruginosa,* respectively (Table 2).

**Table 2.**
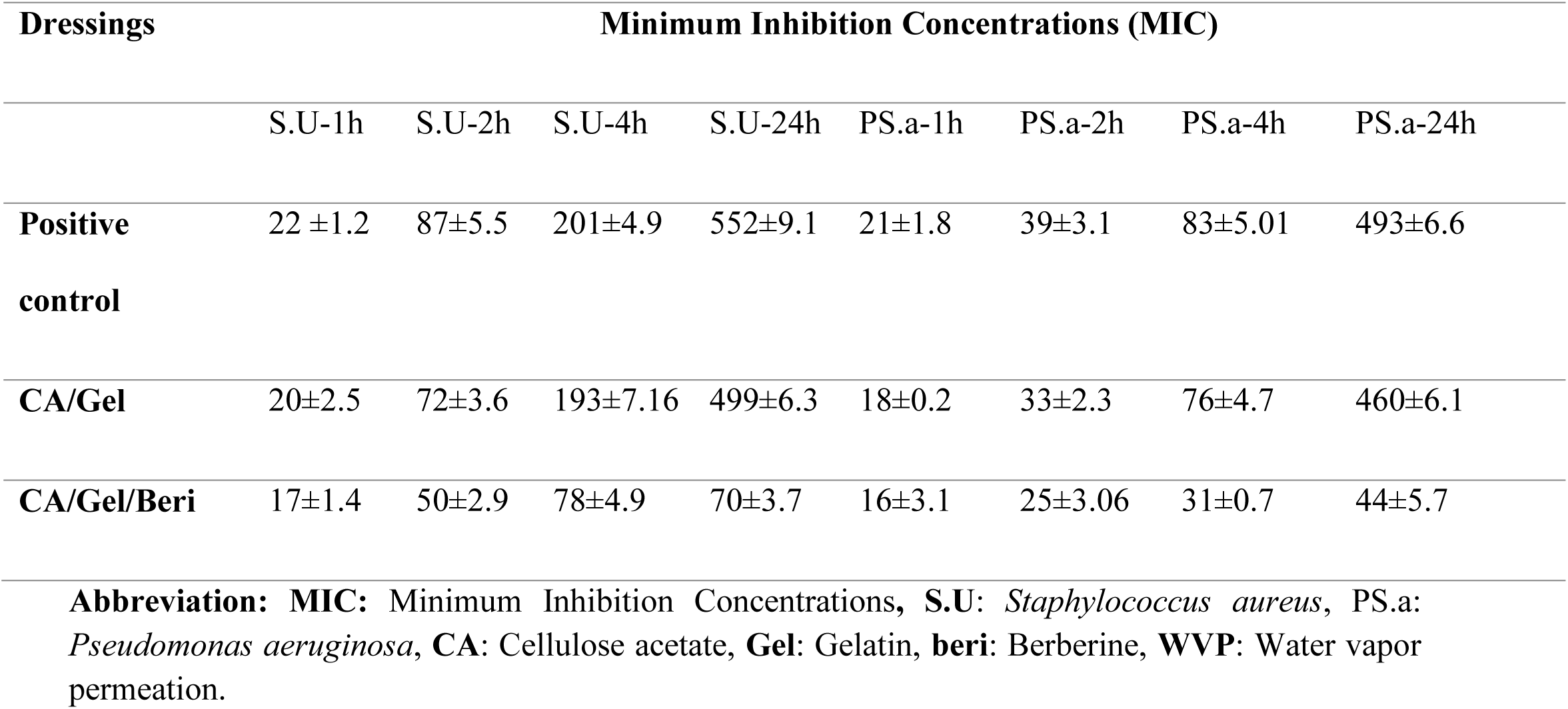
The antibacterial activities of the dressings evaluated by the time-kill assay (number of colony forming unit).

| Dressings | Minimum Inhibition Concentrations (MIC) |  |  |  |  |  |  |  |
| --- | --- | --- | --- | --- | --- | --- | --- | --- |
|  | S.U-1h | S.U-2h | S.U-4h | S.U-24h | PS.a-1h | PS.a-2h | PS.a-4h | PS.a-24h |
| **Positive control** | 22 ±1.2 | 87±5.5 | 201±4.9 | 552±9.1 | 21±1.8 | 39±3.1 | 83±5.01 | 493±6.6 |
| **CA/Gel** | 20±2.5 | 72±3.6 | 193±7.16 | 499±6.3 | 18±0.2 | 33±2.3 | 76±4.7 | 460±6.1 |
| **CA/Gel/Beri** | 17±1.4 | 50±2.9 | 78±4.9 | 70±3.7 | 16±3.1 | 25±3.06 | 31±0.7 | 44±5.7 |

**Abbreviation:** MIC: Minimum Inhibition Concentrations, S.U: *Staphylococcus aureus*, PS.a: *Pseudomonas aeruginosa*, CA: Cellulose acetate, Gel: Gelatin, beri: Berberine, WVP: Water vapor permeation.

The results exhibited that the antibacterial activity of the fabricated CA/Gel/Beri dressing was significantly higher than the positive control and the CA/Gel dressing (p<0.005). Previous studies confirmed the potential antibacterial activity of berberine against various bacteria [46, 48, 49]. Kang et al. [48] reported that berberine exhibited time and concentration dependence antibacterial activity against *Actinobacillus pleuropneumoniae* with the MIC of 0.3125 mg/mL. Wojtyczka et al [50] reported that berberine has antibacterial activity against coagulase-negative staphylococcus strains with the MIC ranged from 16 to 512 μg/mL. The higher bactericidal effect of CA/Gel/Beri dressing observed in our study is related to the incorporation berberine in the matrix of CA/Gel nanofibers, while this value is effective for wound dressing applications. These results imply that the fabricated dressings are not only a barrier against bacterial penetration but also an antibacterial dressing.

### 3.3. The hemocompatibility results

Since wound dressings are designed to contact with bloody wounds, it is essential to assess the hemocompatibility of the fabricated dressings. The hemolysis value induced by the fabricated was measured as an indication of hemocompatibility and the obtained results are presented in Fig. 5.

**Fig. 5.**
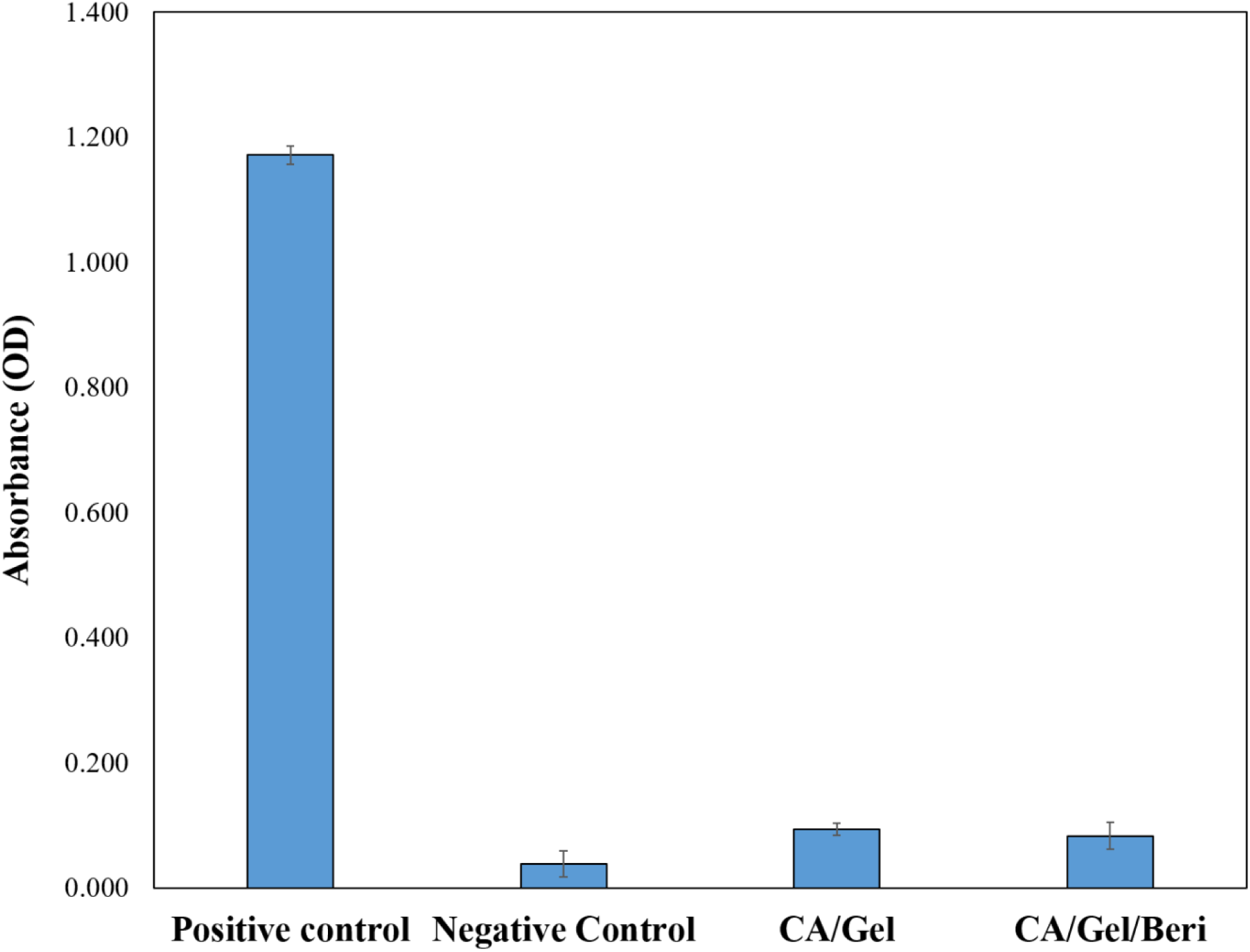
Hemocompatibility histogram of the fabricated dressing. Values represent the mean ± SD, n = 3.

The obtained results from the hemocompatibility test showed that the hemolysis induced by the fabricated dressings were significantly lower than the positive control (distilled water-lysed RBC). Moreover, it is shown that the berberine incorporated dressing exhibited lower hemolysis than pure CA/Gel dressing.

### 3.4. Cell proliferation assay results

The proliferation of L929 murine fibroblastic cell line was measured by MTT assay at 24 and 72 h defter cell seeding and the results are present at Fig. 6.

**Fig. 6.**
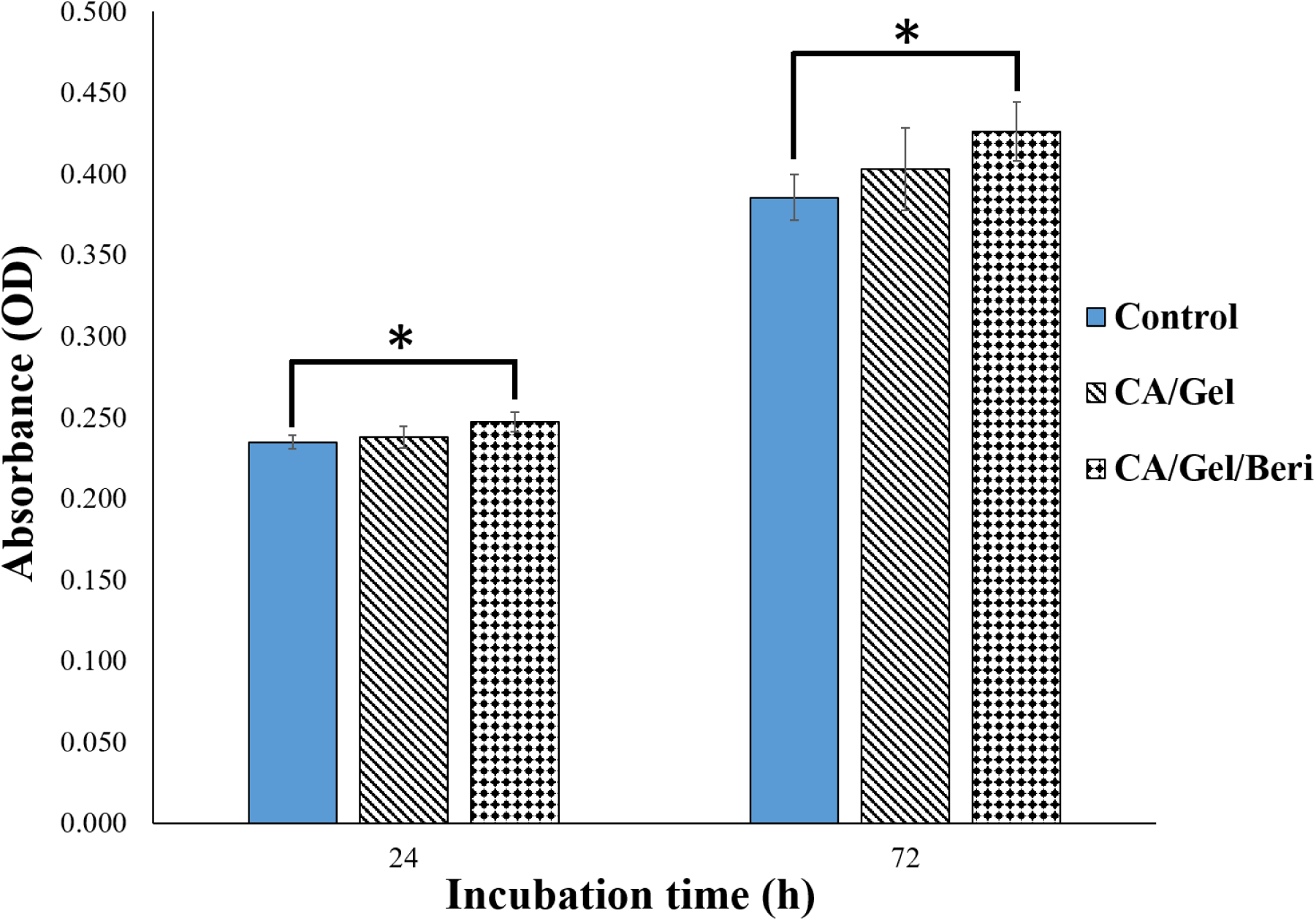
Cell proliferation assay histogram measured by MTT assay after 24 and 72 h cell seeding. Values represent the mean ± SD, n = 3, *: p<0.005.

As shown in Fig. 6, the maximum cell growth was obtained with the CA/Gel/Beri dressing which was statistically significant compared with the control group (tissue culture plate) (p<0.005). It is also apparent that the incorporation of berberine enhanced the proliferation of L929 murine fibroblastic cells. Liu et al. [51] reported that berberine is able to promote the proliferation of periodontal ligament stem cells (hPDLSC) through the activation of ERK-FOS signaling pathway via binding to the EGFR on the cell membrane.

### 3.5. The morphology of the cells on the nanofibers

The morphology of the seeded L929 murine fibroblastic cell on the CA/Gel/Beri was observed using SEM after fixation and dehydration. As shown in Fig. 7, the cultured cells well spread onto the nanofibers after 24 h cell seeding. This image clearly depicts that the cells are attached and spread onto the fabricated nanofibers.

**Fig. 7.**
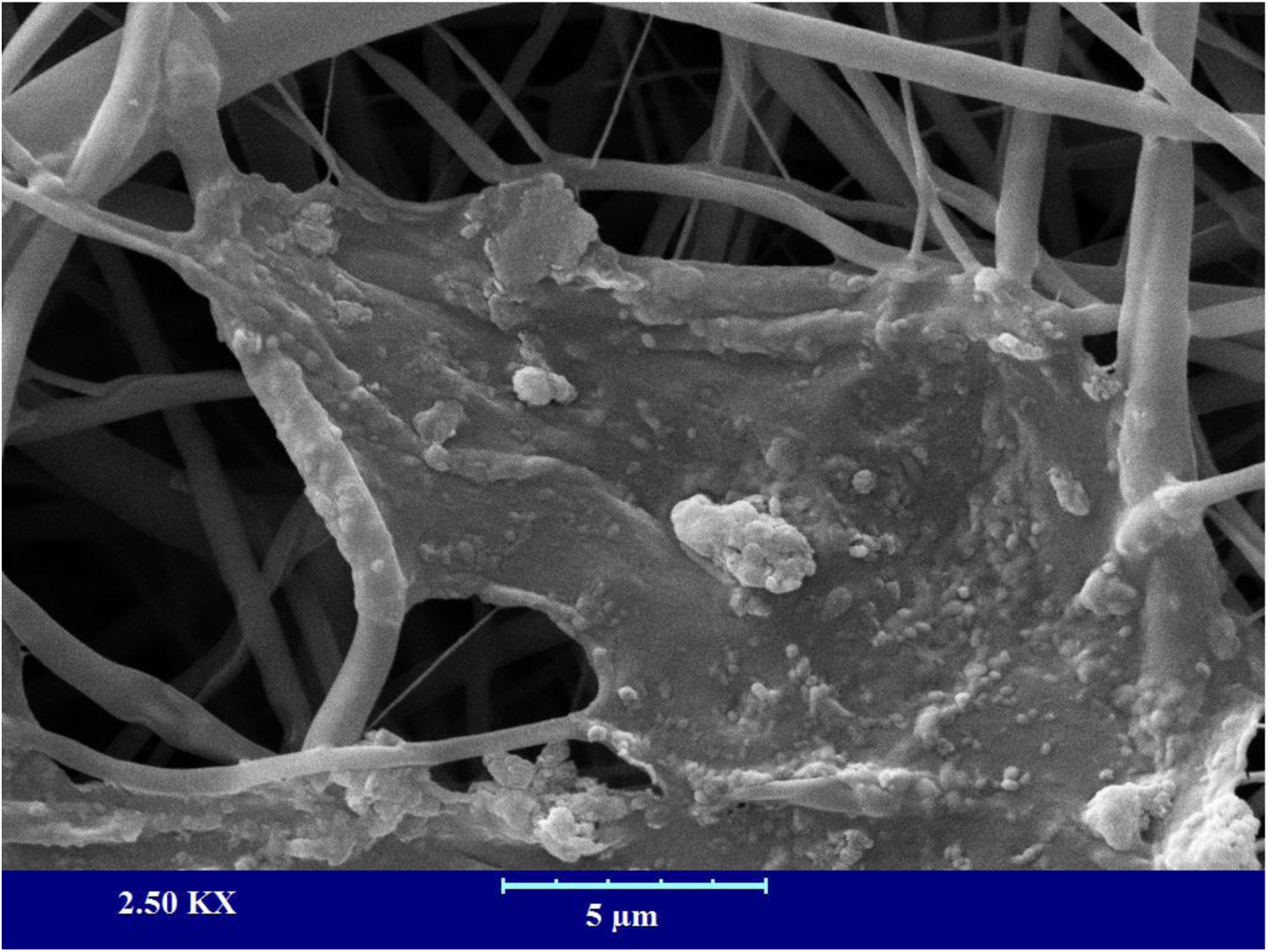
The SEM micrograph of the cultured L929 murine fibroblastic cell on nanofibers after 24 h cell seeding.

### 3.6. Animal study findings

#### 3.6.1. Histopathology and Histomorphometry analysis results

The wound healing efficacy of the fabricated CA/Gel and CA/Gel/Beri dressings was evaluated by the histopathology examinations and the results are presented in Fig. 8. The obtained tissues were stained with H&E and MT staining. Fig. 8a shows the positive control (normal skin) with intact dermal and epidermal, whereas the negative control group (untreated wound) exhibited polymorphonuclear inflammatory cells (PMNs) infiltration and granulation tissue formation, however, the epidermal layer has not been formed yet (Fig. 8b).

**Fig. 8.**
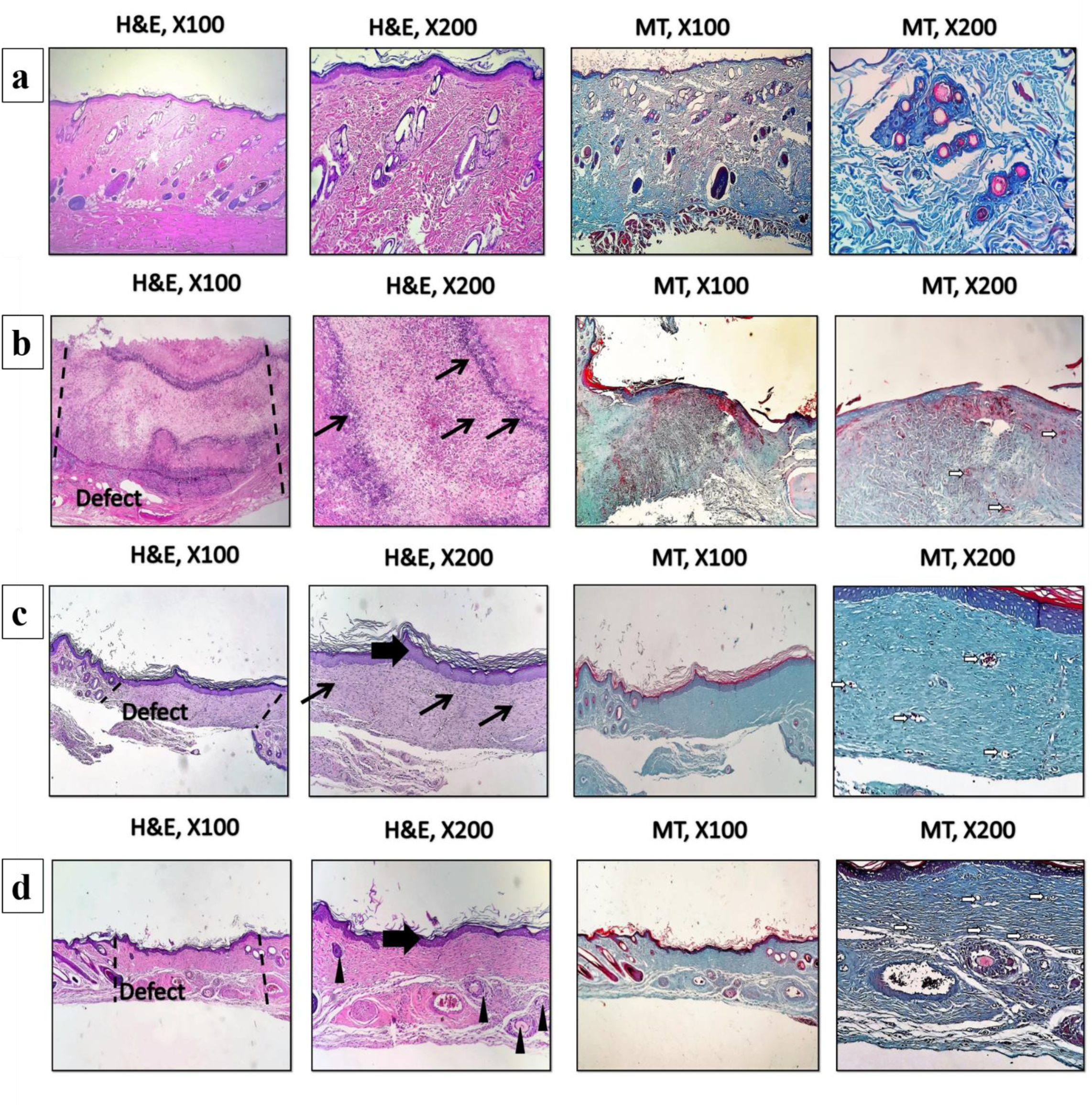
Haematoxylin and Eosin (H&E) and Masson’s trichrome (MT) stained microscopic sections of wounded tissue treated with dressings. (a) The positive control, (b) the negative control, (c) the CA/Gel dressing, and (d) the CA/Gel/Beri dressing. Thick arrows: epidermal layer, thin arrows: infiltration of inflammatory cells, arrowheads rejuvenation of skin appendages, white arrows: neo-vascularization.

The CA/Gel dressing group showed the completed epithelialization and lower infiltrated PMNs in comparison with the negative control group (Fig. 8c). The anti-inflammatory activity of berberine has been shown in other studies. Xiao et al. [52] reported that berberine has an anti-inflammatory effect through the adjusting PCSK9-LDLR pathway and reducing the plasma concentration of IFNγ, TNFα, IL-1α, and 8-isoprostane. Yi et al. [53] demonstrated that berberine acts as an antiatherosclerotic agent through the inhibition of COX-2 expression via the ERK1/2 signaling pathway. In another study, Kim et al. [54] conducted a study to assess the anti-inflammatory effect of berberine on normal human keratinocytes. They reported that berberine treatment reduced IL-6 expression, activity, and expression of matrix metalloproteinase-9 (MMP-9). Moreover, they observed that berberine suppressed AP-1 DNA binding activity and ERK activation.

The epidermal proliferation and the epidermal layer enlargement were evident in CA/Gel/Beri group, while the inflammatory response and the granulation tissue decreased in this group. Rejuvenation of skin appendix was seen and developed in CA/Gel/Beri group. Moreover, this group exhibited more resemblance to normal skin with a thin epidermis, the presence of normal rete ridges, and normal thickness of skin layers. The histopathological observations showed that the fabricated CA/Gel/Beri dressing resulted in the best outcome compared to negative control and CA/Gel groups.

The histomorphometric analysis was done to further evaluate the healing process of the wounded skin (Table 3). The minimum re-epithelialization was observed in the negative control group, while the CA/Gel/Beri exhibited the highest re-epithelialization after 16 days (P< 0.05). The wound site in the negative control group was filled with immature granulation tissue which indicated the slow and ineffective wound healing process.

**Table 3.**
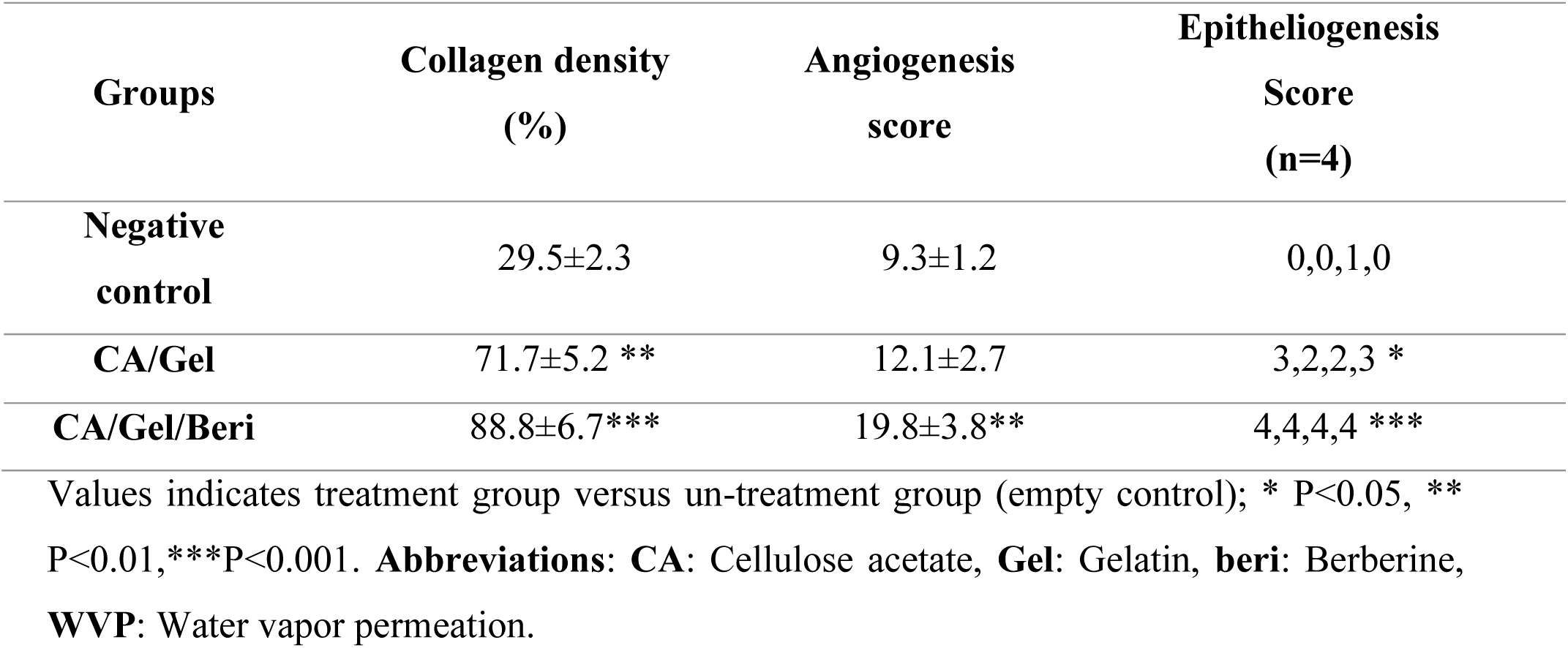
Histomorphometric analysis of different experimental groups.

The wound healing process is critically dependent on the angiogenesis as well as the collagen synthesis in the wound site. In this regard, the angiogenesis and the collagen synthesis were measured to further assess the wound healing process under the influence of the fabricated dressings. The highest collagen density, 88.8±6.7 %, was observed in CA/Gel/Beri which was statistically significant compared with the CA/Gel, 71.7±5.2 %, and the negative control group, 29.5±2.3 %, (P<0.01 and P<0.001). Moreover, CA/Gel/Beri exhibited the highest angiogenesis, 19.8±3.8, while the lowest angiogenesis, 9.3±1.2, was observed in the negative control group (P<0.001). Previous studies demonstrated the wound healing potential of berberine treatment. Pashaee et al. [37] evaluated the wound healing activity of berberine on STZ-induced diabetic rats and conducted that the treatment improved the wound healing process. They proposed that the observed healing process is related to the anti-diabetic, anti-inflammatory, as well as antimicrobial effects of berberine. Our findings indicated that the incorporation of berberine into the dressing significantly improved the healing process.

## Conclusion

DFU is one of the main disabling complications of mellitus diabetes which requires excessive consideration and wound care management. The healing process in diabetic foot ulcer is delayed and take a long time to heal, so an accelerating agents should be applied to enhance the healing process. Moreover, the wound site must be isolated from pathogenic microorganisms via a well-designed dressing. Electrospun nanofibrous dressings are ideal wound dressing due to their nanometric scale, high surface to volume ratio, adjustable porosity, and ability to load various drugs and bioactive molecules. In the present study CA/Gel nanofibrous dressing containing berberine was fabricated using the electrospinning technique and after characterization used as a DFU dressing material. The characterization results demonstrated that the fabricated dressing was suitable as the wound dressing material and did not induced any adverse effect on the cultured cells and also exhibited antibacterial activity against gram-positive and gram-negative bacterium. The animal studies on STZ-induced diabetic rat clearly demonstrated that the CA/Gel/Beri dressing enhanced wound healing process. The evidence from this study suggests that the fabricated CA/Gel/Beri dressing is suitable dressing material to enhance the healing process of diabetic foot ulcer and also isolate the wound site from the pathogenic microorganisms invasion.

## Acknowledgment

The authors gratefully acknowledge the research council of Kermanshah University of Medical Sciences (grant no. 95322) for financial support.

## Authors’ contributions

All authors read and approved the final manuscript.

## References

1. Wynn, M., The efficacy of negative pressure wound therapy for diabetic foot ulcers: A systematised review. Journal of Tissue Viability, 2019.

2. Dow, C., et al., Diet and risk of diabetic retinopathy: A systematic review. European journal of epidemiology, 2018. 33(2): p. 141–156.

3. Jeffcoate, W.J., et al., Current challenges and opportunities in the prevention and management of diabetic foot ulcers. Diabetes care, 2018. 41(4): p. 645–652.

4. Moura, L.I., et al., Recent advances on the development of wound dressings for diabetic foot ulcer treatment—a review. Acta biomaterialia, 2013. 9(7): p. 7093–7114.

5. Mulder, G., D.G. Armstrong, and S. Seaman, Standard, appropriate, and advanced care and medical-legal considerations: Part one-diabetic foot ulcerations. Wounds, 2003. 15(4): p. 92–106.

6. Jannesari, M., et al., Composite poly (vinyl alcohol)/poly (vinyl acetate) electrospun nanofibrous mats as a novel wound dressing matrix for controlled release of drugs. International journal of nanomedicine, 2011. 6: p. 993.

7. Tamer, T., et al., MitoQ loaded chitosan-hyaluronan composite membranes for wound healing. Materials, 2018. 11(4): p. 569.

8. Dhivya, S., V.V. Padma, and E. Santhini, Wound dressings–a review. BioMedicine, 2015. 5(4).

9. Maver, T., et al., Functional wound dressing materials with highly tunable drug release properties. RSC Advances, 2015. 5(95): p. 77873–77884.

10. Samadian, H., S.S. Zakariaee, and R. Faridi-Majidi, Evaluation of effective needleless electrospinning parameters controlling polyacrylonitrile nanofibers diameter via modeling artificial neural networks. The Journal of The Textile Institute, 2018: p. 1–10.

11. Khoshnevisan, K., et al., Cellulose acetate electrospun nanofibers for drug delivery systems: Applications and recent advances. Carbohydrate polymers, 2018.

12. Samadian, H., et al. Needleless electrospinning system, an efficient platform to fabricate carbon nanofibers. in Journal of Nano Research. 2017. Trans Tech Publ.

13. Massoumi, B., et al., Novel nanostructured star-shaped polythiophene, and its electrospun nanofibers with gelatin. Journal of Polymer Research, 2016. 23(7): p. 136.

14. Mirjalili, M. and S. Zohoori, Review for application of electrospinning and electrospun nanofibers technology in textile industry. Journal of Nanostructure in Chemistry, 2016. 6(3): p. 207–213.

15. Hassiba, A.J., et al., Review of recent research on biomedical applications of electrospun polymer nanofibers for improved wound healing. Nanomedicine, 2016. 11(6): p. 715–737.

16. Zhu, Y., C. Romain, and C.K. Williams, Sustainable polymers from renewable resources. Nature, 2016. 540(7633): p. 354.

17. Abbasian, M., et al., Scaffolding polymeric biomaterials: Are naturally occurring biological macromolecules more appropriate for tissue engineering? International Journal of Biological Macromolecules, 2019.

18. Kubo, R., T. Saito, and A. Isogai, Preparation and characterization of carboxylated cellulose nanofibrils with dual metal counterions. Cellulose: p. 1–11.

19. Fukui, S., et al., Surface-hydrophobized TEMPO-nanocellulose/rubber composite films prepared in heterogeneous and homogeneous systems. Cellulose, 2019. 26(1): p. 463–473.

20. Shimizu, M., et al., Thermal and electrical properties of nanocellulose films with different interfibrillar structures of alkyl ammonium carboxylates. Cellulose, 2019: p. 1–9.

21. Hondo, H., T. Saito, and A. Isogai, Preparation of oxidized celluloses in a TEMPO/NaBr system using different chlorine reagents in water. Cellulose, 2019. 26(5): p. 3021–3030.

22. Choong, F.X., et al., Stereochemical identification of glucans by a donor–acceptor–donor conjugated pentamer enables multi-carbohydrate anatomical mapping in plant tissues. Cellulose: p. 1–12.

23. Yaich, A.I., U. Edlund, and A.-C. Albertsson, Barriers from wood hydrolysate/quaternized cellulose polyelectrolyte complexes. Cellulose, 2015. 22(3): p. 1977–1991.

24. Arca, H.C., et al., Synthesis and characterization of alkyl cellulose ω-carboxyesters for amorphous solid dispersion. Cellulose, 2017. 24(2): p. 609–625.

25. Taepaiboon, P., U. Rungsardthong, and P. Supaphol, Vitamin-loaded electrospun cellulose acetate nanofiber mats as transdermal and dermal therapeutic agents of vitamin A acid and vitamin E. European Journal of Pharmaceutics and Biopharmaceutics, 2007. 67(2): p. 387–397.

26. Tungprapa, S., I. Jangchud, and P. Supaphol, Release characteristics of four model drugs from drug-loaded electrospun cellulose acetate fiber mats. Polymer, 2007. 48(17): p. 5030–5041.

27. Gouma, P., et al., Nano-hydroxyapatite—Cellulose acetate composites for growing of bone cells. Materials Science and Engineering: C, 2012. 32(3): p. 607–612.

28. Pishnamazi, M., et al., Microcrystalline cellulose, lactose and lignin blends: Process mapping of dry granulation via roll compaction. Powder Technology, 2019. 341: p. 38–50.

29. Liu, X., et al., Antimicrobial electrospun nanofibers of cellulose acetate and polyester urethane composite for wound dressing. Journal of Biomedical Materials Research Part B: Applied Biomaterials, 2012. 100(6): p. 1556–1565.

30. Yue, K., et al., Synthesis, properties, and biomedical applications of gelatin methacryloyl (GelMA) hydrogels. Biomaterials, 2015. 73: p. 254–271.

31. Kuijpers, A.J., et al., Cross-linking and characterisation of gelatin matrices for biomedical applications. Journal of Biomaterials Science, Polymer Edition, 2000. 11(3): p. 225–243.

32. Massoumi, B., et al., Surface functionalization of graphene oxide with poly (2-hydroxyethyl methacrylate)-graft-poly (ε-caprolactone) and its electrospun nanofibers with gelatin. Applied Physics A, 2016. 122(12): p. 1000.

33. Li, M., et al., Electrospinning polyaniline-contained gelatin nanofibers for tissue engineering applications. Biomaterials, 2006. 27(13): p. 2705–2715.

34. Huang, Y., et al., In vitro characterization of chitosan–gelatin scaffolds for tissue engineering. Biomaterials, 2005. 26(36): p. 7616–7627.

35. García, A., et al., Stability and rheological study of sodium carboxymethyl cellulose and alginate suspensions as binders for lithium ion batteries. Journal of Applied Polymer Science, 2018. 135(17): p. 46217.

36. Gulfraz, M., et al., Comparison of the antidiabetic activity of Berberis lyceum root extract and berberine in alloxan-induced diabetic rats. Phytotherapy Research, 2008. 22(9): p. 1208–1212.

37. Pashaee, M., The effect of hydroalcoholic extract of Berberis vulgaris on wound healing of diabetic wistar rats. Journal of Chemical Health Risks, 2016. 6(4).

38. Ehterami, A., et al., Chitosan/alginate hydrogels containing Alpha-tocopherol for wound healing in rat model. Journal of Drug Delivery Science and Technology, 2019.

39. Farzamfar, S., et al., Taurine-loaded poly (ε-caprolactone)/gelatin electrospun mat as a potential wound dressing material: In vitro and in vivo evaluation. Journal of Bioactive and Compatible Polymers, 2018. 33(3): p. 282–294.

40. Salehi, M., et al., Preparation of pure PLLA, pure chitosan, and PLLA/chitosan blend porous tissue engineering scaffolds by thermally induced phase separation method and evaluation of the corresponding mechanical and biological properties. International Journal of Polymeric Materials and Polymeric Biomaterials, 2015. 64(13): p. 675–682.

41. Semyari, H., et al., Fabrication and characterization of collagen–hydroxyapatite-based composite scaffolds containing doxycycline via freeze-casting method for bone tissue engineering. Journal of biomaterials applications, 2018. 33(4): p. 501–513.

42. Ai, A., et al., Sciatic nerve regeneration with collagen type I hydrogel containing chitosan nanoparticle loaded by insulin. International Journal of Polymeric Materials and Polymeric Biomaterials, 2018: p. 1–10.

43. Farzamfar, S., et al., Neural tissue regeneration by a gabapentin-loaded cellulose acetate/gelatin wet-electrospun scaffold. Cellulose, 2018. 25(2): p. 1229–1238.

44. Kandhare, A.D., P. Ghosh, and S.L. Bodhankar, Naringin, a flavanone glycoside, promotes angiogenesis and inhibits endothelial apoptosis through modulation of inflammatory and growth factor expression in diabetic foot ulcer in rats. Chemico-biological interactions, 2014. 219: p. 101–112.

45. Liu, Y., et al., Tunable physical properties of ethylcellulose/gelatin composite nanofibers by electrospinning. Journal of agricultural and food chemistry, 2018. 66(8): p. 1907–1915.

46. Peng, L., et al., Antibacterial activity and mechanism of berberine against Streptococcus agalactiae. International journal of clinical and experimental pathology, 2015. 8(5): p. 5217.

47. Purna, S.K. and M. Babu, Collagen based dressings--a review. Burns: journal of the International Society for Burn Injuries, 2000. 26(1): p. 54.

48. Kang, S., et al., The antibacterial mechanism of berberine against Actinobacillus pleuropneumoniae. Natural product research, 2015. 29(23): p. 2203–2206.

49. Sarkar, A.K., S. Appidi, and A.S. Ranganath, Evaluation of berberine chloride as a new antibacterial agent against gram-positive bacteria for medical textiles. Fibres & Textiles in Eastern Europe, 2011. 19(4): p. 131–134.

50. Wojtyczka, R., et al., Berberine enhances the antibacterial activity of selected antibiotics against coagulase-negative Staphylococcus strains in vitro. Molecules, 2014. 19(5): p. 6583–6596.

51. Liu, J., et al., The promotion function of berberine for osteogenic differentiation of human periodontal ligament stem cells via ERK-FOS pathway mediated by EGFR. Scientific reports, 2018. 8(1): p. 2848.

52. Xiao, H.-B., et al., Berberine inhibits dyslipidemia in C57BL/6 mice with lipopolysaccharide induced inflammation. Pharmacological Reports, 2012. 64(4): p. 889–895.

53. Yi, G., et al., Biochemical pathways in the antiatherosclerotic effect of berberine. Chinese medical journal, 2008. 121(13): p. 1197–1203.

54. Kim, S., et al., Berberine inhibits TPA-induced MMP-9 and IL-6 expression in normal human keratinocytes. Phytomedicine, 2008. 15(5): p. 340–347.

